# Nucleation of Biomolecular Condensates from Finite-Sized Simulations

**DOI:** 10.1101/2022.11.11.515961

**Authors:** Lunna Li, Matteo Paloni, Aaron R. Finney, Alessandro Barducci, Matteo Salvalaglio

## Abstract

The nucleation of protein condensates is a concentration-driven process of assembly. When modelled in the canonical ensemble, condensation is affected by finite-size effects. Here, we present a general and efficient route to obtain ensemble properties of protein condensates in the macroscopic limit from finite-sized nucleation simulations. The approach is based on a theoretical description of droplet nucleation in the canonical ensemble and enables estimating thermodynamic and kinetic parameters, such as the macroscopic equilibrium density of the dilute protein phase, the condensates surface tension and nucleation free energy barriers. We apply the method to coarse-grained simulations of NDDX4 and FUS-LC, two phase-separating disordered proteins with different physicochemical characteristics. Our results show that NDDX4 condensate droplets, characterised by lower surface tension, higher solubility, and faster monomer exchange dynamics than FUS-LC, form with negligible nucleation barriers. In contrast, FUS-LC condensates form via an activated process over a wide range of concentrations.

Graphical Table of Contents.

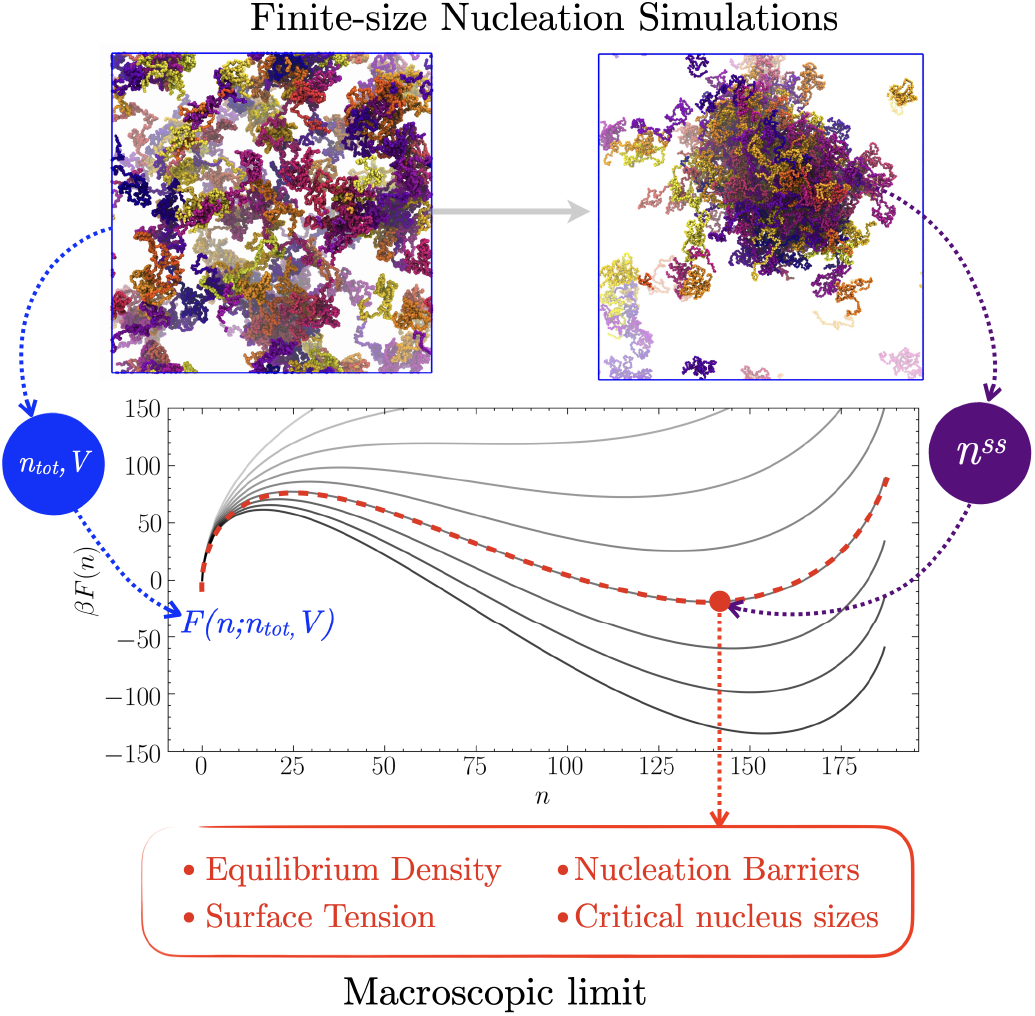

---

Biomolecular compartments that are not bound by membranes have attracted a lot of attention in the last decade because of their important role in cellular organization. ^1,2^ The assembly of these membraneless organelles (MLOs) is driven by the formation of dynamical multivalent interactions between proteins and/or nucleic acids, ^1,2^ often following a nucleation mechanism. ^3,4^

Notably, many proteins involved in forming such compartments are either intrinsically disordered or have highly-flexible domains: typical examples include proteins from the DEAD-box ^5–7^ and FET ^8–10^ families. The interactions between biomacromolecules in the formation of MLOs was described according to the stickers-and-spacers framework, derived for associative polymers. ^9,11^ In this framework, polymer chains are characterized by multivalent domains or motifs, named stickers, that govern intermolecular interactions, interspersed by spacer domains that influence the material properties of the condensates. ^9^ Experiments of simplified systems with one to a few of these disordered protein regions have shown that they can lead to the formation of assemblies via a process of liquid-liquid phase separation (LLPS). ^5,6,8^ Still, the phenomenon in cells could be more complex, involving different molecular mechanisms. ^12–14^

Molecular simulations of simplified systems have provided important insights into the mechanisms and molecular drivers for the formation of biomolecular assemblies. Explicit-solvent molecular dynamics (MD) simulations offer a detailed picture of the structure of the intermolecular interactions and of the relationship between local structure and phase separation propensity. ^15–18^ Still, system sizes and time scales that can be investigated using atomistic MD are severely limited. To alleviate these difficulties, several Coarse-Grained (CG) models with a one-bead-per-residue resolution were proposed ^19–23^ and successfully applied to study phase separating systems, establish coexistence conditions, and the effect of mutations and post-translational modifications on phase separation. ^19,23,24^ Unfortunately, even CG simulations suffer from size limitations and particular strategies, such as the slab method, ^19^ have to be adopted for minimizing finite-size effects in the study of phase-separation processes. Indeed, in finite-sized systems, the free-energy change associated with the assembly of a condensate droplet is a function of both the concentration and the total volume of the system. ^25–29^ This dependence emerges from the fact that, in small volumes, ^30^ the chemical potential of the environment surrounding a condensate droplet depends on its size, leading to qualitative and quantitative differences compared to its macroscopic counterpart. ^26,29,31^ An elegant approach to account for finite-size effects is the Modified Liquid Droplet (MLD) model, ^25,26^ which provides an expression for the nucleation free energy *F* (*n*) in the canonical ensemble under the same set of assumptions typically adopted by classical nucleation theory (CNT). Expressed as a function of the density of the dilute (*ρ_d_*) and condensed (*ρ_c_*) phases, the MLD *F* (*n*) reads:

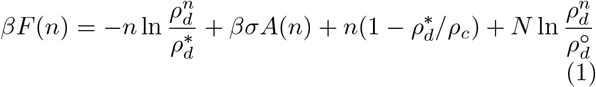

where *β* = 1*/kT*, *k* is the Boltzmann constant, *T* is the temperature, *N* is the total number of chains contained in a simulation box of volume *V*, *n* is the number of chains in the condensate droplet, *σ* is the planar surface tension of the condensed phase, *A*(*n*) is the surface area of a droplet of condensed phase formed by *n* chains, *ρ_c_* is the equilibrium molar density of the condensed phase, 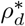 is the equilibrium molar density of the dilute phase, 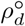 is the total protein density *N/V*, and 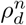 is the density of the dilute phase in a system where *n* chains form a condensed phase droplet at constant *N* and *V* :

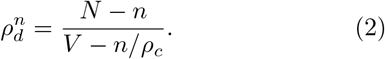

Examples of *F* (*n*) profiles for systems of increasing size are represented in Fig. 1E, where the stationary points correspond to the critical nucleus size *n*^∗^ and to the size of the self-limiting steady-state droplet *n_ss_*, the values of which both increase in magnitude as the volume and the number of molecules in the system increases. For any concentration, one can always identify a threshold volume below which condensation is inhibited by finite-size effects since *F* (*n*), becomes a monotonically increasing function of droplet size, *n*. ^25–29^

**Figure 1:**
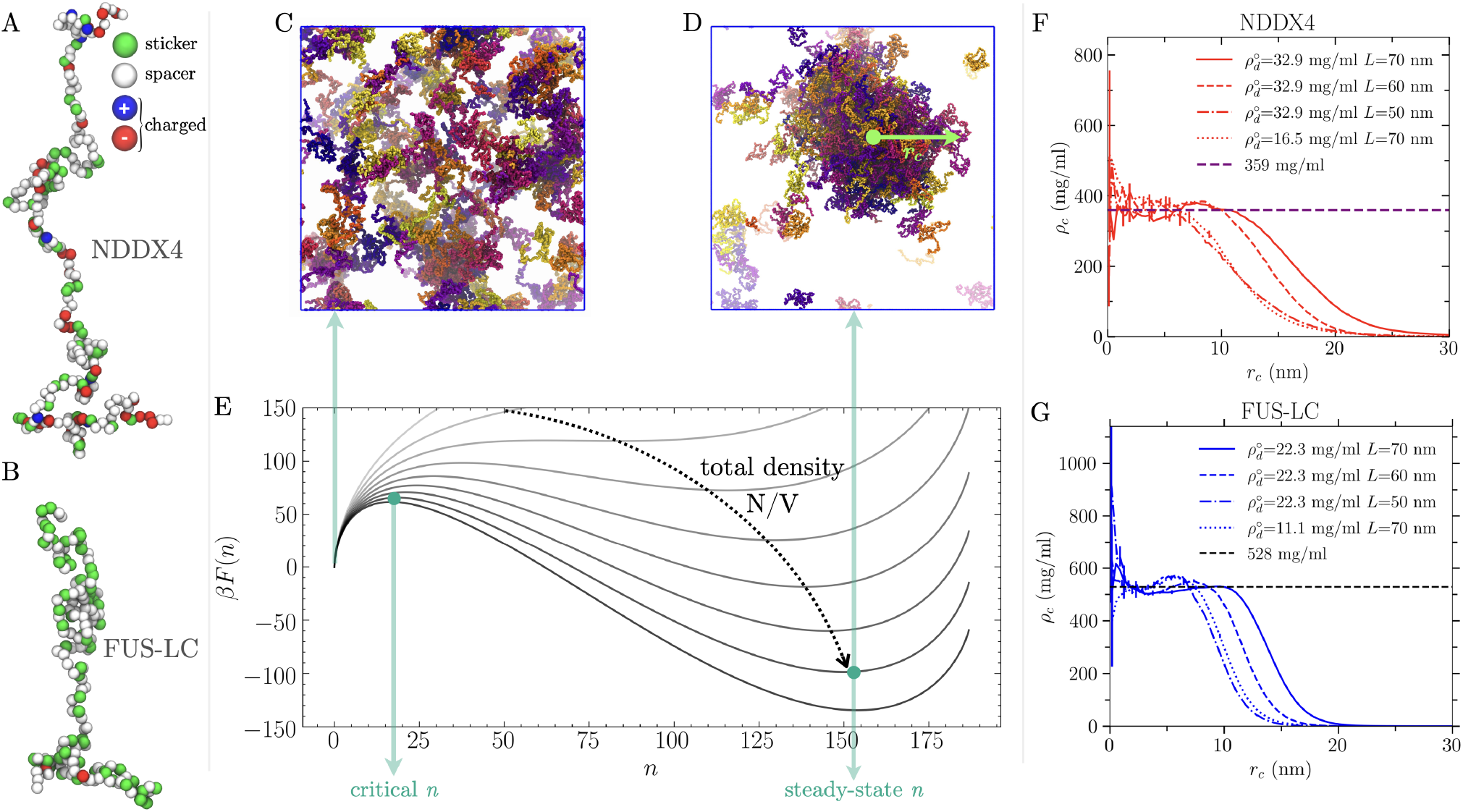
Coarse-grained modelling of the nucleation of bio-molecular condensates in the *NV T* ensemble. **A-B)** Coarse-grained model of the NDDX4 and FUS-LC chains. **C)** Example of supersaturated homogeneous dilute phase configuration. **D)** Example of a steady-state configuration containing a stable, condensed phase droplet. **E**) Example free energy profiles associated with the nucleation of condensed phase droplets, obtained by keeping the total number of peptide chains constant while increasing the total volume. The variable *n* represents the number of chains in the dense phase; the origin corresponds to a homogeneous dilute phase; the local maximum at small *n* corresponds to the critical nucleus size; and the local minimum at large *n* corresponds to the steady-state droplet size. The free energies are obtained using from Eq. 1 with *σ*=0.182 mN/m, 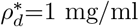, a total number of chains of 187, and a box-size ranging from 105 to 68 nm. **F, G)** Steady-state droplet radial density profiles obtained from simulations performed in different box sizes for NDDX4 (**F**) and FUS-LC (**G**).

Here we adopt this theoretical framework to fully account for the finite-size effects described above to characterize the thermodynamics and kinetics of biomolecular condensation in MD simulations with tractable system sizes. We demonstrate the potential of our approach by focusing on two intrinsically-disordered protein domains: FUS-LC and NDDX4, which are popular model systems for investigating biomolecular condensates. ^6,9^ While both proteins undergo LLPS at ambient temperature, they display markedly distinct physicochemical characteristics. Notably, FUS-LC is enriched in aromatic and/or polar residues Gln and Ser ^9^ whereas the NDDX4 sequence is relatively abundant in charged residues organized in patches of opposite signs, significantly contributing to its condensation behavior. ^6^

We model these proteins using a sequence-specific CG model with a one-bead-per-residue resolution. Consecutive amino acids are connected using a harmonic potential with an equilibrium distance of 0.38 nm and a spring constant of 1 × 10^3^ kJ/mol/nm^2^. Non-bonded amino acids interact through electrostatic interactions and short-range contact potentials. Electrostatic interactions between charged amino acids are described with a Debye-Hückel potential with a screening length of 1 nm, corresponding approximately to an ionic strength of 100 mM. Short-range non-bonded interactions are modelled using a Lennard-Jones potential with residue-dependent *σ* and *ϵ* (see Tab. S2) inspired by the stickers-spacers framework. ^9,32^ In this respect, we take advantage of mutagenesis studies that suggested a key role of amino-acids with large-sized aromatic or planar side chains in driving the phase separation of our model systems. ^6,9,33^ Thus we define Arg, Phe, Tyr, Trp and Gln residues as Stickers (St) and all the others as Spacers (Sp), and we set the LJ potential as 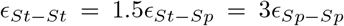. The absolute energy scale of these short-range interactions is the only free parameter of our model and we tuned it to reproduce the experimental densities of NDDX4 and FUS-LC proteins. ^9,33,34^ Functional forms of the interactions and parameters of the CG model are available in SI.

Using this CG potential, we performed a large set of NVT simulations of NDDX4 or FUS-LC to probe condensation from homogeneous solutions at diverse densities and sizes of the systems in standard cubic boxes (see Table 1). For all the systems, we performed simulations for 1 *μs* to equilibrate the systems, followed by an additional 1 *μs* for analyses of the resulting steady state. All the simulations were conducted at a temperature of 300 K and an ionic strength of 100 mM in GROMACS 2019.4 (see the SI for further details). We simulated supersaturation regimes sufficient to allow condensed phase formation within a reasonable simulation time, while avoiding extremely high concentrations that may result in large condensates spanning across simulation box periodic boundaries.

**Table 1:** Modified-liquid-droplet nucleation simulations

| System | $L(\text{nm})$ | $N$ | $\rho_d^\circ$<br>($\text{nm}^{-3}$) | $\rho_d^\circ$<br>($\text{mg/ml}$) | $n_{ss}$ |
| --- | --- | --- | --- | --- | --- |
| NDDX4-1 | 40 | 25 | 0.00039 | 16.5 | × |
| NDDX4-2 | 40 | 35 | 0.00055 | 23.2 | △ |
| NDDX4-3 | 40 | 50 | 0.00078 | 32.9 | △ |
| NDDX4-4 | 50 | 49 | 0.00039 | 16.5 | △ |
| NDDX4-5 | 50 | 69 | 0.00055 | 23.2 | 45(2) |
| NDDX4-6 | 50 | 98 | 0.00078 | 32.9 | 76(2) |
| NDDX4-7 | 60 | 84 | 0.00039 | 16.5 | 42(2) |
| NDDX4-8 | 60 | 118 | 0.00055 | 23.2 | 84(3) |
| NDDX4-9 | 60 | 169 | 0.00078 | 32.9 | 139(2) |
| NDDX4-10 | 70 | 133 | 0.00039 | 16.5 | 71(2) |
| NDDX4-11 | 70 | 187 | 0.00055 | 23.2 | 130(3) |
| NDDX4-12 | 70 | 268 | 0.00078 | 32.9 | 214(2) |
| FUS-LC-1 | 40 | 25 | 0.00039 | 11.1 | × |
| FUS-LC-2 | 40 | 35 | 0.00055 | 15.7 | △ |
| FUS-LC-3 | 40 | 50 | 0.00078 | 22.3 | 42(0.5) |
| FUS-LC-4 | 50 | 49 | 0.00039 | 11.1 | △ |
| FUS-LC-5 | 50 | 69 | 0.00055 | 15.7 | 55(1) |
| FUS-LC-6 | 50 | 98 | 0.00078 | 22.3 | 86(1) |
| FUS-LC-7 | 60 | 84 | 0.00039 | 11.1 | 63(1) |
| FUS-LC-8 | 60 | 118 | 0.00055 | 15.7 | 101(1) |
| FUS-LC-9 | 60 | 169 | 0.00078 | 22.3 | 153(1) |
| FUS-LC-10 | 70 | 133 | 0.00039 | 11.1 | 103(2) |
| FUS-LC-11 | 70 | 187 | 0.00055 | 15.7 | 158(1) |
| FUS-LC-12 | 70 | 268 | 0.00078 | 22.3 | 243(2) |

$n_{ss}$ is the average steady-state cluster size with block average error in the bracket; △ indicates fluctuating cluster (unstable primary droplet/presence of multiple droplets) or that the box sizes are too small to distinguish dilute/dense phase; × indicates no phase separation.

In the majority of simulations, small droplet condensates of stationary size *n_ss_* - corresponding to local minima in Fig. 1E - were observed. We identified droplets according to a two-dimensional geometric criterion, considering the protein inter-chain contacts and distances between the centres of mass (COM) of individual chains and the droplet COM (see Supporting Information). Analysis of protein radial number density profiles from the COM of steady-state droplets (see Figs. 1 F, G) indicate that densities in the core are independent of the size and overall concentration of the simulated system. The mean densities in this region provide a robust estimate of the equilibrium dense-phase densities for NDDX4 (359 mg/ml) and FUS-LC (527 mg/ml), which are in good agreement with values obtained by slab coexistence simulations (NDDX4: 336 mg/ml, FUS-LC: 484 mg/ml) and experimental results (NDDX4: 380 mg/ml, ^33^ FUS-LC: 477 mg/ml ^34^) (see SI, Coexistence Simulations section). Conversely, the direct estimate of dilute-phase density from finite-volume nucleation simulations can suffer from artefacts that result in errors that can scale as *V* ^1*/*4^. ^35^

We, therefore, rely on the MLD framework that, for finite-sized systems, provides the following Gibbs-Thomson/Kelvin equation for the equilibrium density of the dilute phase: ^25^

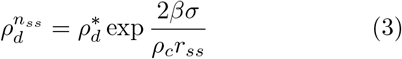

where 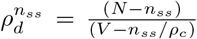, and *r_ss_* is the radius of the steady-state droplet. Under the assumption of spherical droplets, one can explicitly introduce the dependence of the coexistence pressure and the droplet radius *r_ss_* on the number of chains in the droplet and reformulate Eq. 3 as:

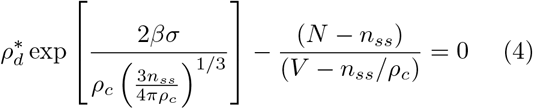

This equation provides the opportunity to fully characterize the assembly process by determining all the key thermodynamic quantities from our finite-size simulations. Indeed, in Eq. 4 *N*, *V* and *T* are defined by the simulation setup while *ρ_c_* and *n_ss_* can be directly obtained from the analysis of the droplet density profiles (see Figs. 1F, G; and additional details reported in SI). Most importantly, the dilute phase density 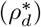 and the surface tension (*σ*) can be computed by a global fitting of Eq.4 on the data from our set of simulations performed at different values of *N* and *V*. Using this strategy, we estimated 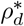 to be 1.03 ± 0.32 mg/ml for FUS and 4.87 ± 1.44 mg/ml for NDDX4 (error bars indicate 95% confidence intervals from bootstrapped results). Both densities are in excellent agreement with estimates obtained using slab coexistence simulations based on the same CG model: 1.42 ± 0.30 mg/ml and 4.78 ± 0.88 mg/ml for FUS-LC and NDDX4, respectively. Both the slab and nucleation results are comparable to the experimental dilute-phase values of 2 mg/ml ^9^ and 7 mg/ml. ^33^ The surface tension *σ* is estimated to be 0.37 ± 0.11 mN/m, for FUS-LC and 0.101 ± 0.06 mN/m for NDDX4, reflecting the higher hydrophilic character of NDDX4 compared with FUS-LC. These estimates are in excellent agreement with surface tension estimates computed from slab simulations, ^36^ yielding 0.125 ± 0.099 and 0.291 ± 0.026 mN/m for NDDX4 and FUS-LC, respectively. Notably, our FUS-LC surface tension estimate agrees with calculations performed by Benayad et al. ^37^ for an explicit solvent FUS-LC coarse grained model, which identified the surface tension in the 0.01–0.4 mN/m range from fluctuations of the droplet shape and from the broadening of the interface between phases.

To validate the equilibrium density and surface tension parameters obtained from fitting simulation data with Eq. 4, we compare the position of the minima in the function *F* (*n*) (Eq. 1), parameterised with 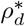 and *σ* obtained from fitting, with the steady-state droplet size measured in simulations (see Fig. S5). The excellent agreement shown by the parity line demonstrates the method’s power for universal calculations of thermodynamic properties such as surface tension and equilibrium vapour pressure using finite-sized nucleation simulations.

Importantly, the determination of 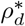 and *σ* (as well as *ρ_c_*) enables the calculation of free energy profiles in the limit of an infinitely large simulation box. This is achieved by evaluating Eq. 1 in the limit where *N* ≫ *n*, and *V* ≫ *nv_ℓ_*. In this limit, 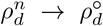, and Eq. 1 reduces to the typical CNT expression for reversible work of formation of a condensate droplet from a supersaturated dilute phase at constant density 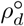:

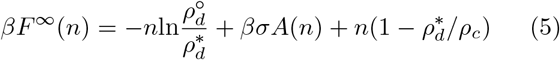

Using this approach, we can thus predict nucleation free energy barriers and critical nucleus sizes under conditions that should be optimal for comparison with experiments without the need for computationally-demanding schemes able to mimic open-boundary/infinite-reservoir macroscopic conditions. ^38,39^

Fig. 2 demonstrates how the proposed methodology provides a direct route to extract the macroscopic thermodynamic parameters that enable such characterisation from simple finite-sized simulations. Examples of finite-size free energy profiles obtained for NDDX4 and FUS-LC, are provided in Fig. 2A at one of the total densities simulated. Nucleation free energies in the macroscopic limit are instead reported for all densities investigated in Fig. 2B. From the nucleation free energy profiles, we can obtain nucleation barrier estimates and therefore discuss relative nucleation kinetics in the limit of a macroscopic open system. For instance, the nucleation barriers for FUS-LC are consistently higher and are associated with larger critical nuclei cf. NDDX4 (see Fig. 2B) under the conditions studied. We note, however, that using a CNT-based model for nucleation with thermodynamic parameters evaluated using our computational approach indicates a cross-over in the nucleation free energy barrier (and thus in the nucleation rates) for densities lower than those explicitly simulated. This can be seen in Fig. 2D, where the nucleation free energy barrier for FUS-LC becomes lower than that of NDDX4 below 10 mg/mL. The critical nuclei sizes also display a crossover, observed at higher density values of approximately 16 mg/mL. The difference in the crossover density between free energy barrier height and critical nucleus size reflects the differences in the balance between surface and bulk free energy terms for NDDX4 and FUS-LC.

**Figure 2:**
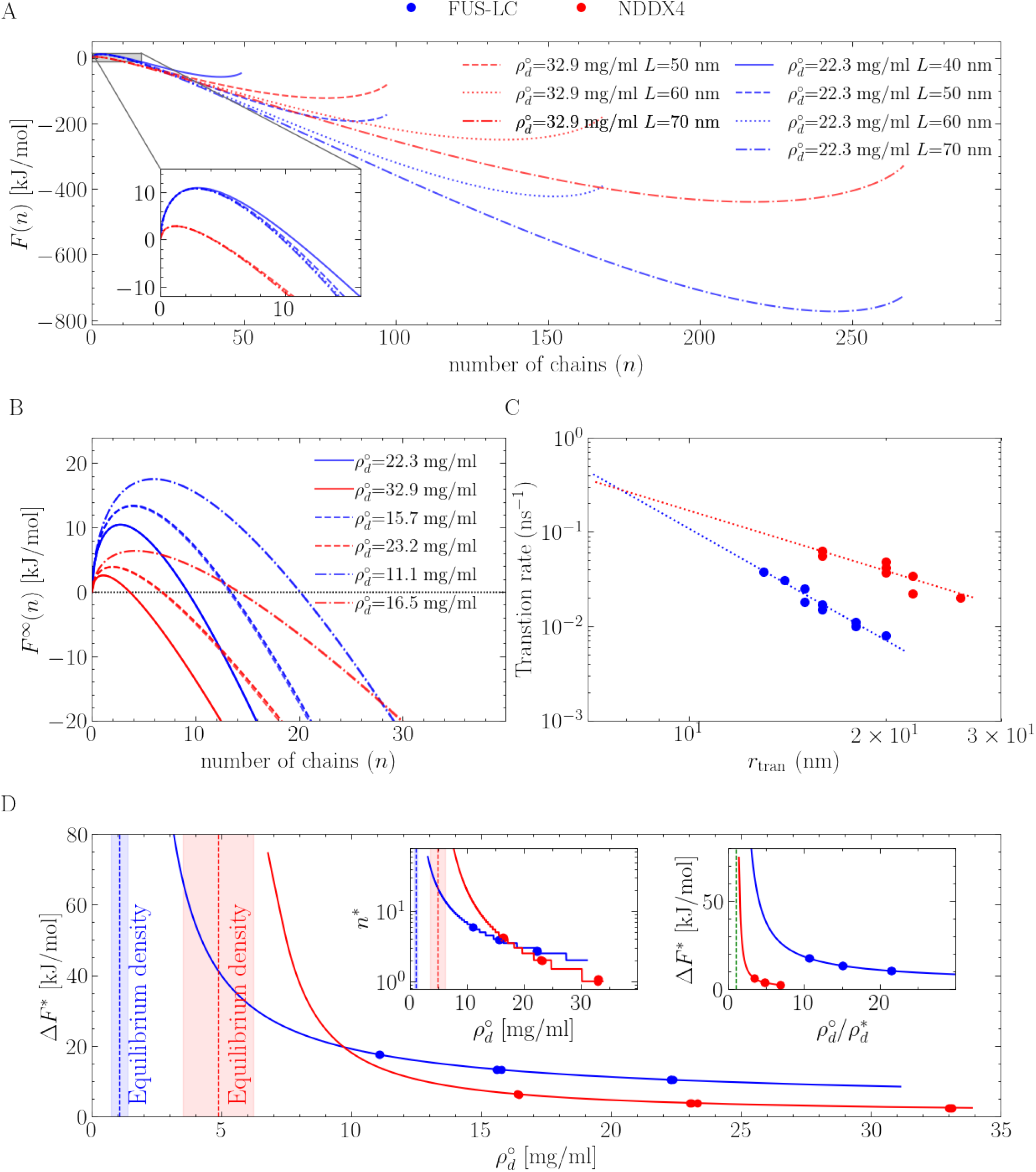
Protein condensate nucleation simulations for NDDX4 (red) and FUS-LC (blue). **A)** Free energy profiles associated with the nucleation of condensed phase droplets. The variable *n* represents the number of chains in the condensed phase. **B)** Macroscopic free energy profiles, with the effect of the artificial confinement associated with *NV T* simulations removed. For the each protein, curves of increasing barriers correspond to reducing bulk densities/supersaturation. **C)** The rate of condensed-to-dilute transition for droplets of different sizes and the exponential fit. The rate is calculated as the reverse of the mean-first-passage-time of the condensed-to-dilute transition from a Markov-state model (see Figs. S7 and S8). **D)** Estimates of free energy barriers and critical nucleus sizes (left inset) at different bulk densities/supersaturation. The equilibrium density of the dilute phase is reported as a dashed line, with the shaded area representing the 95% confidence interval computed from a bootstrap analysis. The right inset of **D)** is a representation of **D)** plotted in supersaturation.

Differences in the physicochemical character of NDDX4 and FUS-LC are further reflected by their different solubility (captured by 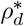), which induces different supersaturation levels at the same bulk densities.

Approaching the binodal line (the green dashed line in the right inset of Fig. 2D, corresponding to 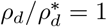), both the critical nucleus size and Δ*F*^∗^ diverge. At these conditions, directly observing nucleation events is extremely unlikely, even in the limit of very large simulations. ^31^ As such, complete information on the nucleation behavior approaching the binodal can only be inferred from theory.

At steady state, nucleation simulations provide an extensive sampling of the dynamic exchange of monomers between the condensed and dilute phases. We can exploit this to obtain a quantitative description of the single chain exchange dynamics, with the aim of complementing the collective information captured by nucleation free energy profiles. To this aim, we analyse the dynamics of single-chain exchange between the condensed and dilute phases using Markov state models (MSMs). The details of the MSM construction and analysis are reported in Supporting Information. Briefly, in the MSMs, the dilute and condensed states are identified based on the inter-chain contacts and on the distance between the COM of the individual chains and the droplet (see Figs. S7, S8). As a measure of the exchange dynamics, we compute the rate of escape of a single chain from the condensed to the dilute phase as a function of the droplet size (see Table S1, Fig. S8). Fig. 2C shows an approximately exponential decay in the condensed-to-dilute transition rate as a function of droplet size, with FUS-LC demonstrating a steeper decay than NDDX4. Moreover, irrespective of the droplet size, it is slower to transfer a FUS-LC chain across the phase boundary compared with NDDX4. The estimated rate of dilute-to-condensed transitions appears instead largely uncorrelated with respect to the droplet size, but, as expected for a diffusion-dominated process, it fluctuates around the same average for both NDDX4 and FUS-LC. The ratio of the condensed-to-dilute and dilute-to-condensed transition rates is approximately linear with respect to the ratio of the number of peptides in the two phases (see Fig. S6). The faster escape dynamics of single chains from NDDX4 condensate droplets impact fluctuations in their overall size and shape. NDDX4 generally shows slightly larger size fluctuations than FUS-LC (see Table 1), possibly due to a combined effect of longer chain length, different hydrophobicity and faster escape dynamics. In addition, while both NDDX4 and FUS-LC condensate droplets can be effectively approximated as spherical in the theoretical analysis and interpretation of the simulation results, we note that NDDX4 faster exchange dynamics, higher hydrophilicity and more gentle radial density gradients (see Fig. 1, panels F, G) lead to larger deviations from a perfectly spherical shape (see Table S1).

Using the results obtained from nucleation simulations, we can rationalize the effect of finite size on the thermodynamics of phase separation by inserting *σ* and 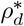 into Eq. 1, thus mapping the qualitative nucleation behavior as a function of the total peptide density and system size. Following this strategy, we produce domain diagrams in Fig. 3 that indicate the presence or absence of phase separation for both NDDX4 and FUS-LC. In the blue/red shaded region, exemplified by state I, the finite-sized thermodynamics admit the existence of steady state droplets corresponding to local minima in the nucleation free energy profiles. Increasing the system size and density in this region results in larger droplets, as demonstrated by the size of the circles used to represent our simulation results. In the white region featuring state III, confinement induces a monotonically increasing free energy curve, ^26,29^ and nucleation will never occur regardless of the simulation time. The blue/red solid line represents the transition boundary, where the free energy curve has a single stationary point corresponding to a flex. This condition is closely approximated by the free energy profile of state Simulations initiated on the boundary line from a pre-formed condensate droplet of size close to the stationary point in the free energy profile will experience negligible driving forces to either grow or dissipate. ^40^ At large system sizes this effect manifests itself as a very slow evolution of the droplet towards the equilibrium state, corresponding to a homogeneous phase with density 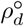. Instead, at small system sizes, where far fewer chains are present, the shallow free energy gradient results in large fluctuations of the droplet size. This behavior is confirmed by simulations performed in regimes of volume and peptide density closely approximating these conditions (see Table 2).

**Table 2:** Dissolution/nucleation simulations at larger system sizes and lower densities

| System | $L(\text{nm})$ | $N$ | $\rho_d^\circ(\text{nm}^{-3})$ | $\rho_d^\circ(\text{mg/ml})$ | Initial condition | Result | Simulation length ($\mu\text{s}$) |
| --- | --- | --- | --- | --- | --- | --- | --- |
| NDDX4-13 | 120 | 169 | 0.00010 | 4.2 | NDDX4-9 | dissolved | 2 |
| NDDX4-14 | 120 | 268 | 0.000155 | 6.5 | NDDX4-12 | dissolved | 4 |
| NDDX4-15 | 174 | 1000 | 0.00019 | 8.0 | homogeneous | no nucleation | 5 |
| NDDX4-16 | 205 | 1000 | 0.00012 | 5.1 | homogeneous | no nucleation | 5 |
| FUS-LC-13 | 120 | 133 | 0.00008 | 2.3 | FUS-10 | dissolved | 1 |
| FUS-LC-14 | 120 | 187 | 0.00011 | 3.1 | FUS-11 | dissolved | 6 |
| FUS-LC-15 | 242 | 1000 | 0.00007 | 2.0 | homogeneous | no nucleation | 5 |
| FUS-LC-16 | 329 | 1000 | 0.00003 | 0.9 | homogeneous | no nucleation | 5 |

**Figure 3:**
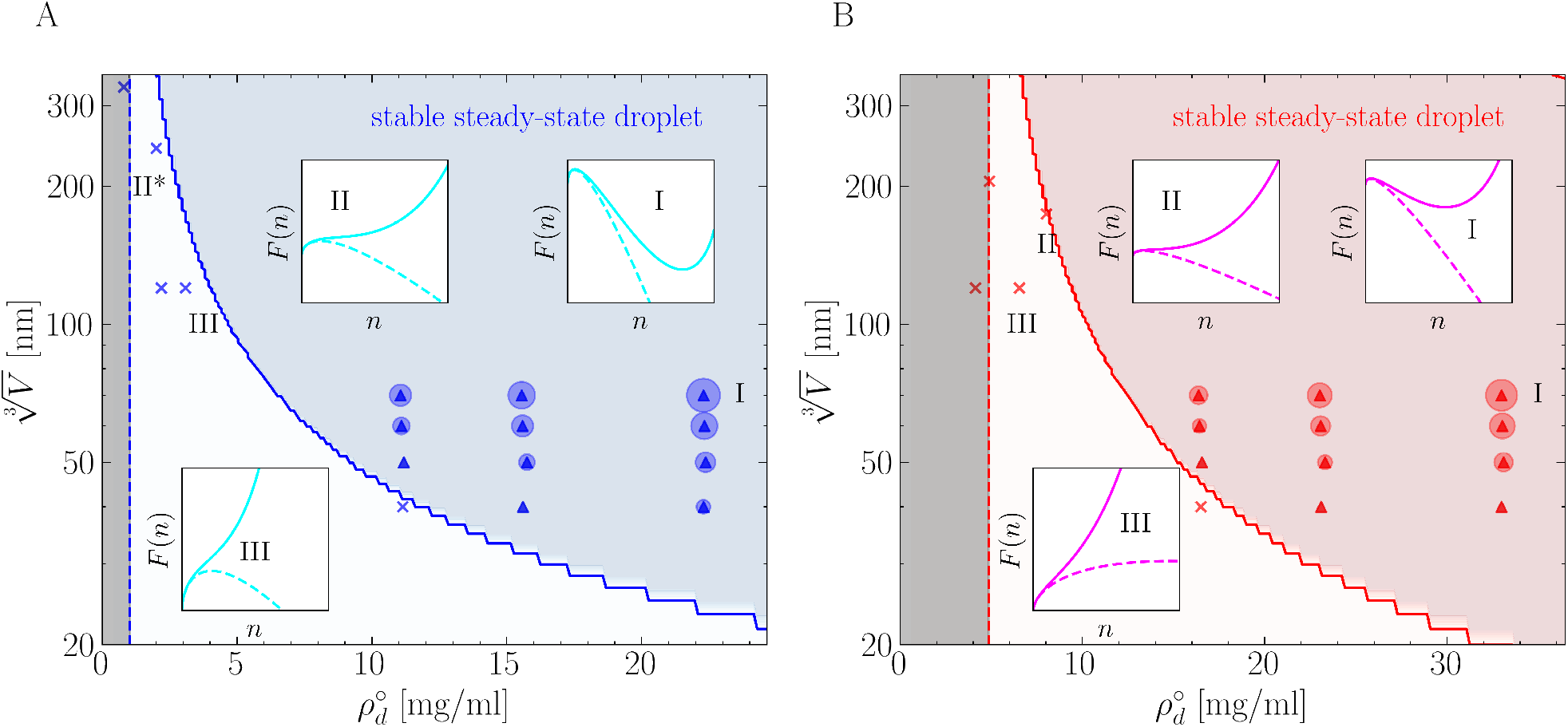
Stability of FUS-LC **A)** and NDDX4 **B)** dense-phase droplets in confined volumes as a function of the total peptide density, predicted using parameters fitted from simulation data from Tables 1 and 2. The grey shaded region represents the region 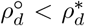 where phase separation is thermodynamically unstable, with the dashed lines representing the predicted equilibrium vapour density 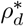. Nucleation is thermodynamically favoured above 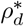. The white region represents a *nominally supersaturated* portion of parameter space; here finite-size effects mean that, *F* (*n*) (represented as a solid line in the insets) is a monotonically increasing function of *n*, even if the corresponding free energy in the macroscopic limit (represented as a dashed line) admits a critical nucleus. In these conditions condensation is inhibited by the confinement (see inset III). Simulations performed in these conditions (indicated with ×) cannot show nucleation or relax to a homogeneous vapour phase when initialised from a droplet. The blue/red shaded region instead represents the ensemble of conditions in which droplets are thermodynamically stable, characterized by the free energy profile of state II. All simulations performed in this region show phase separation. The simulations used to fit the thermodynamic parameters are represented with circles with sizes proportional to the volume of the corresponding steady-state droplets. The blue/red solid line between the white and the blue/red shaded regions is characterized by the free energy profile of boundary state II. All of the free energy profiles in the insets correspond to simulation data points with the same state labels, except for FUS II, which refers to a slightly higher 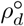 than the actual data point at II*. Inset states of the same numbering are plotted on the same scale for FUS-LC and NDDX4.

Applying a general thermodynamics framework for interpreting nucleation in finite-sized system is a powerful approach to correctly investigate the formation of biomolecular condensates with simulations of reasonable size. This strategy yields consistent estimates of thermodynamic properties of protein condensates, such as the equilibrium density of the dilute and condensed phases and the surface tension. In turn, information on the nucleation kinetics, such as the critical nucleus size and the free energy nucleation barrier can be obtained.

Here we have used this method to quantitatively characterize the LLPS of two model phase-separating systems, such as NDDX4 and FUS-LC, by using simulation based on a minimal, one-bead-per-residue description. Nevertheless, the same approach can be straightforwardly extended to transferrable higher resolution models, ^41,42^ which provide more accurate conformational description and may help rationalizing complex nucleation mechanisms. ^3^ Furthermore, the effect of system size on these thermodynamic and kinetic properties can be clearly demonstrated and quantified. This is particularly relevant in the context of biological systems, where phase separations take place in micrometer-scale isolated cellular compartments, as well as in the rational development of technological applications of LLPS for material synthesis in micro/nanofluidic devices. ^40,43^

## Supporting information

Supplementary Methods and Results

## Acknowledgement

MS and LL gratefully acknowledge the Leverhulme Trust for funding (Project RPG-2019-235). MS and ARF gratefully acknowledge funding from the EPSRC Programme Grant Crystallization in the Real World (Grant EP/R018820/1). AB and MP gratefully acknowledge the support by the French National Research Agency (ANR) under grant ANR-21-CE30-0001 and the Swiss National Science Foundation under the grant CRSII5 193740. The authors acknowledge the use of the UCL High-Performance Computing Facilities and associated support services in the completion of this work.

## Supporting Information Available

Additional details and figures on the clustering algorithms used to identify the number of chains in the dense phase, calculation of the droplet density profiles, Markov State Models for estimating the exchange kinetics, and force-field parameters are available as Supplementary materials. Jupyter notebooks used in the data analyses are available for download at https://github.com/mme-ucl/confined_LLPS.

