## Supplementary Methods and Results for "Nucleation of Biomolecular Condensates from Finite-Sized Simulations"

### Supplementary Material for: Nucleation of Biomolecular Condensates from Finite-Sized Simulations

#### Peptide Sequence

NDDX4:

MGDEDWEAEINPHMSSYVPIFEKDRYSGENGDNFNRTPASSSEMDDGPSRRDHFMKSGFA
SGRNFGNRDAGECNKRDNTSTMGGFGVGKSFGNRGFSNSRFEDGDSSGFWRESSNDCEDN
PTRNRGFSKRGGYRDGNNSEASGPYRRGGRGSFRGCRGGFGLGSPNNDLDPDECMQRTGG
LFGSRRPVLSGTGNGDTSQSRSGSGSERGGYKGLNEEVITGSGKNSWKSEAEGGES

FUS-LC:

MASNDYTQQATQSYGAYPTQPGQGYSQQSSQPYGQQSYSGYSQSTDTSGYGQSSYSSYGQ
SQNTGYGTQSTPQGYGSTGGYGSSQSSQSSYGQQSSYPGYGQQPAPSSTSGSYGSSSQSS
SYGQPQSGSYSQQPSYGGQQQSYGQQQSYNPPQGYGQQNQYNS

### Clustering algorithm

We use PLUMED 2.5.2[1–3] to analyse the steady state condensed phase of NDDX4 and FUS-LC. We adopt a segment-based method in which each peptide chain is represented by different segments. For a NDDX4 molecule of 236 residues, each segment is defined to contain 20 CG beads ($n_{\mathrm{beads}}$=20), with the last segment containing 36 residues. A similar scheme is applied to FUS-LC, which gives a total of 8 segments per FUS-LC chain, with the last segment containing 23 residues. The COM of each segment is then used to construct a radial distribution function (RDF) for the system. Based on the RDF profiles, a threshold distance $R_0$ can be selected to compute the coordination number (inter-chain/intra-chain contact) for the NDDX4/FUS-LC chains using the COORDINDATION function in PLUMED:

$$\sum_{i \in A} \sum_{j \in B} s_{ij} \tag{1}$$

where $s_{ij}$ corresponds to the following switching function:

$$s_{ij} = \frac{1 - (\frac{r_{ij}}{r_0})^6}{1 - (\frac{r_{ij}}{r_0})^{12}} \tag{2}$$

The orange dashed line in Fig. S1 indicates that $R_0$=3.0 nm can be a reasonable choice for NDDX4 and FUS-LC for constructing inter-/intra- chain contacts. Next, the inter- and intra-chain contacts of every peptide chain in the system and over the full simulation length are used to build a 2-dimensional free energy surface (2D-FES). Figure S2 shows that only the inter-chain contact can be used to distinguish different peptide states, as indicated by the dashed blue line. However, it is not sufficient to use one-dimensional (1D) property. As the system size and overall peptide density get smaller, with reduced supersaturation and enhanced finite-size effect, the threshold inter-chain contact tends to shift towards lower values, making it challenging to adapt a universal 1D criteria for distinguish the condensed/dilute phase. Therefore, we also track the distance between the COM of each chain and the condensed phase droplet (COM-distance), to help determining if a peptide chain belong to the dilute or condensed phase. Figures S3 and S4 shows the 2D-FES for cluster analysis of NDDX4 and FUS-LC.

A transition saddle region can be observed in all cases, and it is located at different inter-chain contact/COM-distance values for different system size/peptide density combinations. The orange and cyan dashed lines in Figs. S3, S4 represent the threshold inter-chain contact and COM-distance $r_{\mathrm{tran}}$. For every simulation frame, we assign all the chains with inter-chain contact above the threshold, and with COM-distance below the threshold, as condensed phase chains; all the rest of the chains in the system are considered to be in the dilute phase. Chains that show lower COM-distance but also lower inter-chain contacts represent molecules that are very close to the primary droplet but with much reduced inter-chain interactions; chains with larger COM-distance but moderate inter-chain contact are small clusters of chains in the dilute phase, or a few interface chains that are on the edge of droplet as the condensed phase deviates from a spherical droplet during fluctuation. None are included in the condensed phase counts. For both NDDX4 and FUS-LC, the small clusters of chains prevail in the dilute phase, but any specific cluster typically dissipates within short time, in contrast to the stable condensed phase droplet. Similar phenomena are generally discussed for nucleating events ranging from inorganic compounds[4,5] to biological molecules.[6]

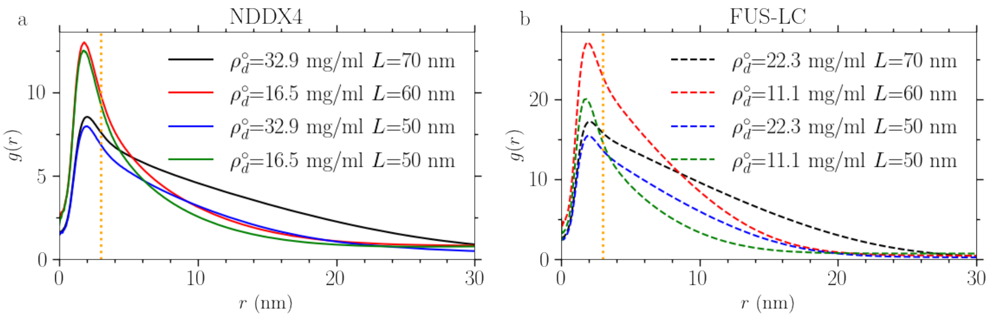


Figure S1: RDF for the segment-COM of NDDX4 and FUS-LC.

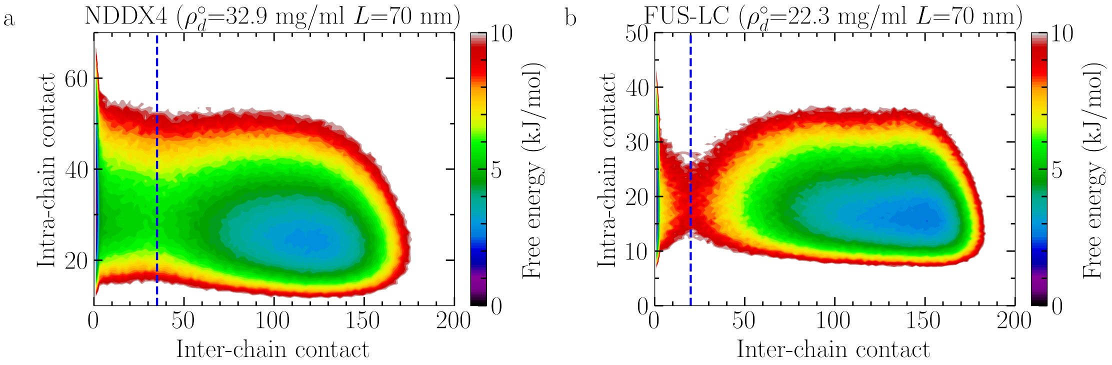


Figure S2: 2D-FES of inter- and intra-chain contacts for NDDX4 and FUS-LC.

Comparison of Figs. S3 and S4 indicates that NDDX4 demonstrates a broader pathway for transition between the dilute and condensed phase. NDDX4 typically shows a larger transition region than FUS-LC on a similar free energy scale. For the limiting case of $\rho_d^\circ = 0.00078$ nm$^{-3}$ and $L = 40$ nm, the NDDX4 droplet is already becoming very fluctuating and much less well-defined due to the combined effect of finite size and reduced supersaturation, while the effect is not yet severe for FUS-LC at the same bulk density/system size condition. For NDDX4 at $\rho_d^\circ = 0.00078$ nm$^{-3}$ and $L = 40$ nm, the threshold COM-distance is $r_{\mathrm{tran}}$=16 nm, comparable to the box size $L/2 = 20$ nm, making it challenging to define the condensed/dilute phase boundary.

Table S1 shows the list of threshold COM-distance $r_{\text{tran}}$ and droplet radius $r_{\text{ss}}$ calculated via $\left(\frac{3n_{ss}v_\ell}{4\pi}\right)^{1/3}$ for NDDX4 and FUS-LC. Droplet size represented by $r_{\text{tran}}$ is consistently larger than $r_{\text{ss}}$, because $r_{\text{tran}}$ also takes into account chains close to the transition saddle region; but $r_{\text{tran}}$ and $r_{\text{ss}}$ are linearly correlated with each other (see Fig. S5 top inset), so they can be both used to depict droplet size.

As discussed in the main text, the predicted $\sigma$ of FUS-LC has smaller errors than NDDX4. The slightly improved accuracy may be reflected in the relative shape anisotropy $\kappa^2$ of the condensed phase droplets between the two peptides. $\kappa^2$ can be used to measures the deviation from a CNT-assumed perfect sphere. Table S1 shows that overall FUS-LC droplet are more spherical than those of NDDX4. As mentioned in the main text, the different sphericity could be associated with the contrasting nature of two peptides, i.e., different hydrophilicity/hydrophobicity. In addition, increased finite-size effect and reduced supersaturation can observe larger deviations from spherical droplet.

Table S1: Droplet size for different system size/bulk combinations

| System | $L$ (nm) | $\rho_d^\circ$ (nm$^{-3}$) | $\rho_d^\circ$ (mg/ml) | $r_{\text{tran}}$ (nm) | $r_{ss} = \left(\frac{3n_{ss}v_\ell}{4\pi}\right)^{1/3}$ (nm) | $\kappa^2$ |
|---|---|---|---|---|---|---|
| FUS-LC-5 | 50 | 0.00055 | 15.7 | 14 | 8.9 | 0.08 |
| FUS-LC-6 | 50 | 0.00078 | 22.3 | 15 | 10.4 | 0.06 |
| FUS-LC-7 | 60 | 0.00039 | 11.1 | 15 | 9.3 | 0.07 |
| FUS-LC-8 | 60 | 0.00055 | 15.7 | 16 | 10.9 | 0.06 |
| FUS-LC-9 | 60 | 0.00078 | 22.3 | 18 | 12.5 | 0.04 |
| FUS-LC-10 | 70 | 0.00039 | 11.1 | 16 | 11 | 0.05 |
| FUS-LC-11 | 70 | 0.00078 | 22.3 | 18 | 12.7 | 0.07 |
| FUS-LC-12 | 70 | 0.00078 | 22.3 | 20 | 14.6 | 0.03 |
| NDDX4-5 | 50 | 0.00055 | 23.2 | 16 | 10.8 | 0.11 |
| NDDX4-6 | 50 | 0.00078 | 32.9 | 20 | 12.9 | 0.15 |
| NDDX4-7 | 60 | 0.00039 | 16.5 | 16 | 10.6 | 0.12 |
| NDDX4-9 | 60 | 0.00055 | 23.2 | 20 | 13.3 | 0.08 |
| NDDX4-9 | 60 | 0.00078 | 32.9 | 22 | 15.7 | 0.06 |
| NDDX4-10 | 70 | 0.00039 | 16.5 | 20 | 12.6 | 0.08 |
| NDDX4-11 | 70 | 0.00055 | 23.2 | 22 | 15.4 | 0.11 |
| NDDX4-12 | 70 | 0.00078 | 32.9 | 26 | 18.2 | 0.05 |

$\kappa^2$ is the relative shape anisotropy[7] computed via MDTraj 1.9.4[8] using COM of chains belong to the condensed phase droplet.

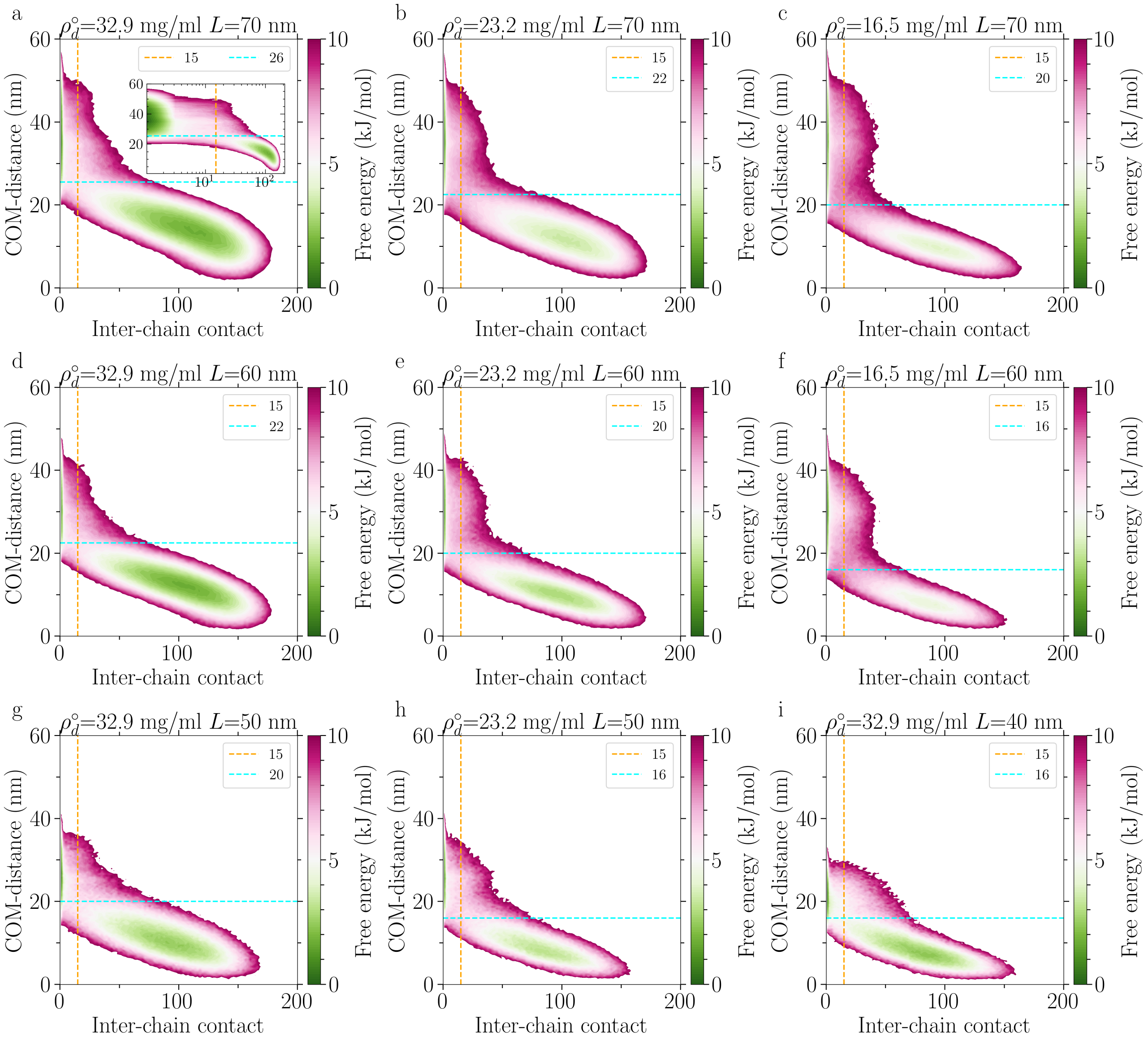


Figure S3: 2D-FES of inter-chain contact and COM-distance for NDDX4

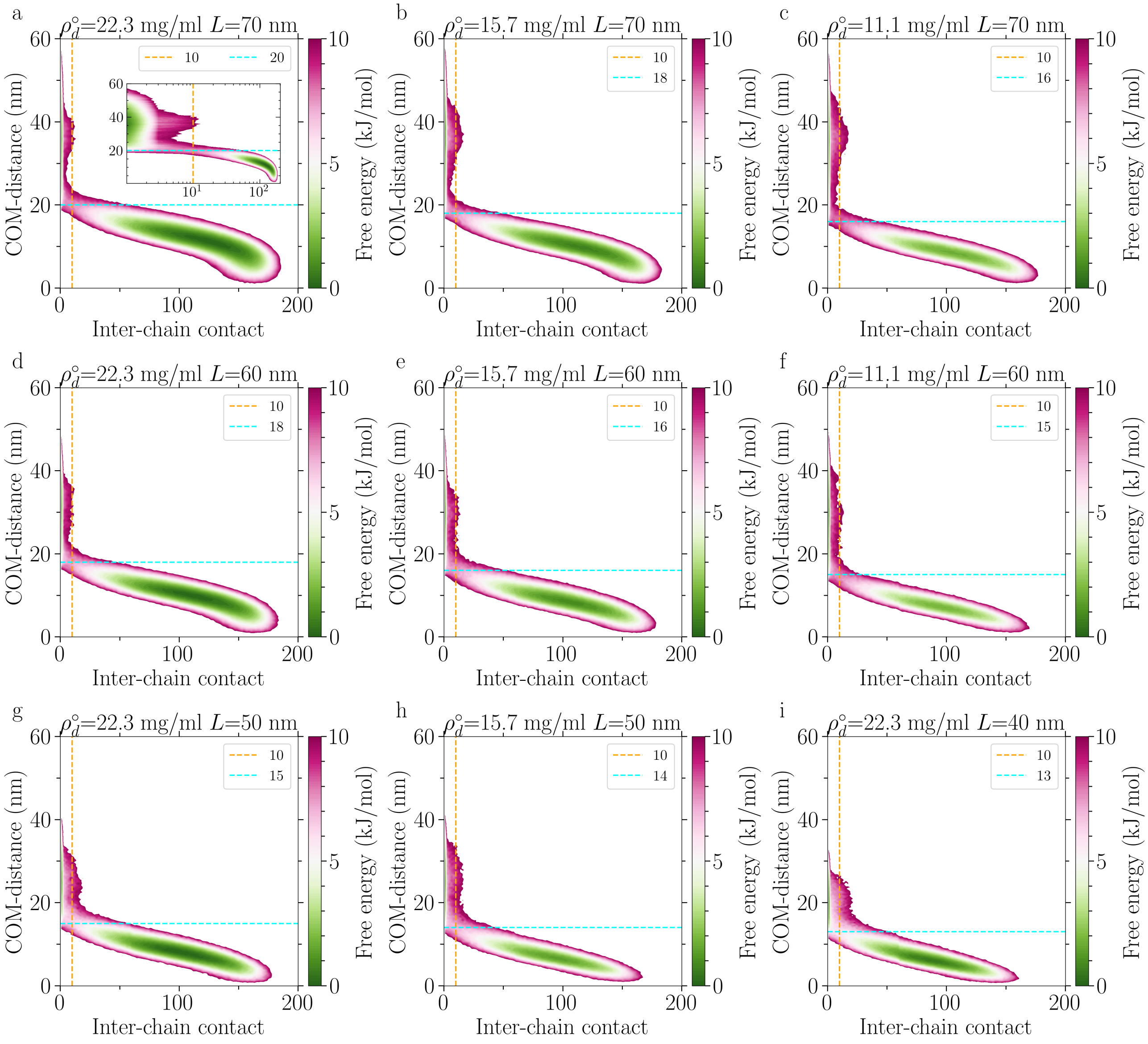


Figure S4: 2D-FES of inter-chain contact and COM-distance for FUS-LC

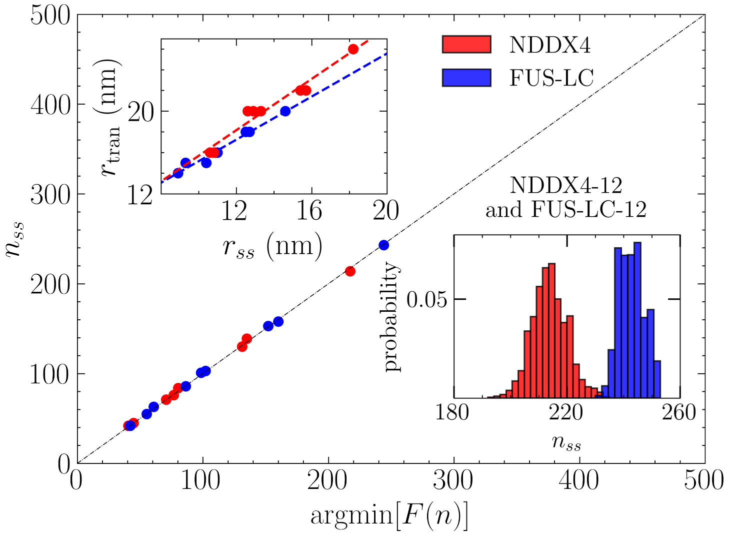


Figure S5: Comparison between the size of the steady-state droplet size measured from simulations and used for performing a global fit of $\sigma$ and $p_d^*$ via Eq. 4 with the position of the local minimum of the free energy profile expressed by Eq. 1. Top inset: comparison of the steady-state droplet radius $r_{ss}$ with the transition radius $r_{\rm tran}$ and the least squares fitted line. Bottom inset: clustersize distribution for NDDX4 at $\rho_d^\circ$=32.9 mg/ml and $L$=70 nm and FUS-LC at $\rho_d^\circ$=22.3 and $L$=70 nm.

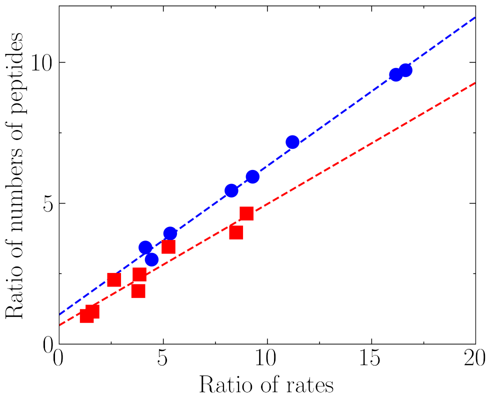


Figure S6: Ratio of the rates of condensed-to-dilute and dilute-to-condensed exchange versus the ratio of the number of peptides in the steady-state condensed and dilute phases for NDDX4 (red) and FUS-LC (blue).

#### Calculation of steady state droplet density profiles

We use the COM of each chain to build the density profiles for FUS-LC and NDDX4 using an in-house analysis package. The COM trajectories in the steady state are used to construct the largest cluster, representing the liquid droplet, based on a minimum distance criterion. Here, the radial distribution functions for FUS-LC and NDDX4 are first computed and the position of the first maximum informed the truncation distance below which two COMs are considered in direct contact. From these contacts, all of the chains in the liquid droplet are identified and the volume number density of the COMs are computed as a function of the radial distance from the centre of mass of the droplet. Density profiles from this analysis are summarized in Figs. 1 F and G. The small number of chains in the core of the droplets result in significant noise in the computed densities in this region, as highlighted by the uncertainty bars in the figures, computed as the standard deviations from block averaging the full trajectory at every $r_c$ windows.

#### Markov-state model

We conduct a 2-dimensional Markov-state model (MSM) analysis[9] using PyEMMA[10] for every system size/bulk pepetide density combinations for NDDX4 and FUS-LC, in order to probe the steady state dynamics of chain exchange between the dilute and condensed phases (see Figs. S7 and S8). Overall, the transition regions obtained from the MSM analysis are similar to those observed from 2D-FES (Figs. S3 and S4). We record the reverse of mean-first-passage-time of condensed-to-dilute transition as the condensed-to-dilute rate for plotting Fig. 2C.

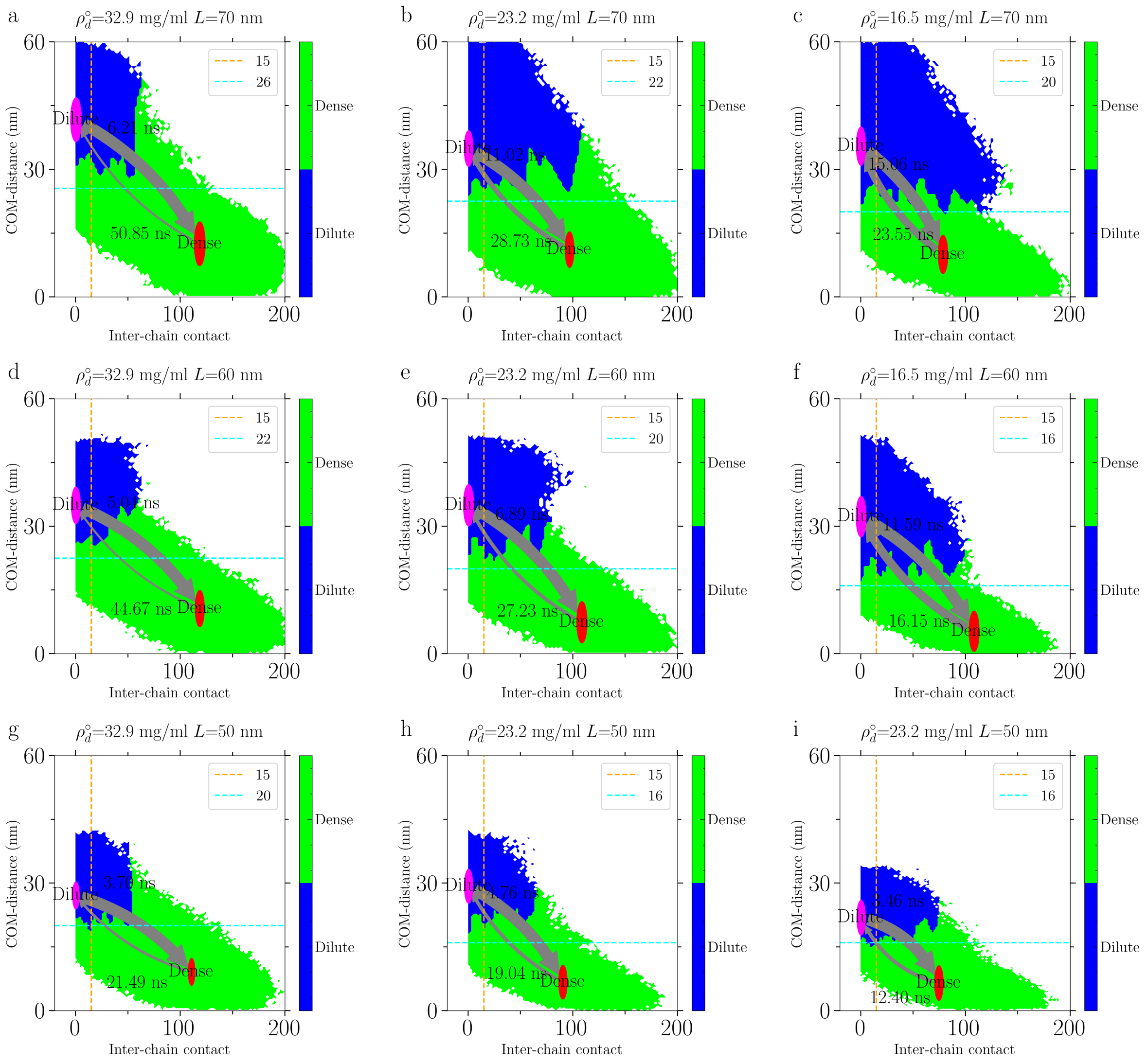


Figure S7: 2D MSM analysis for NDDX4

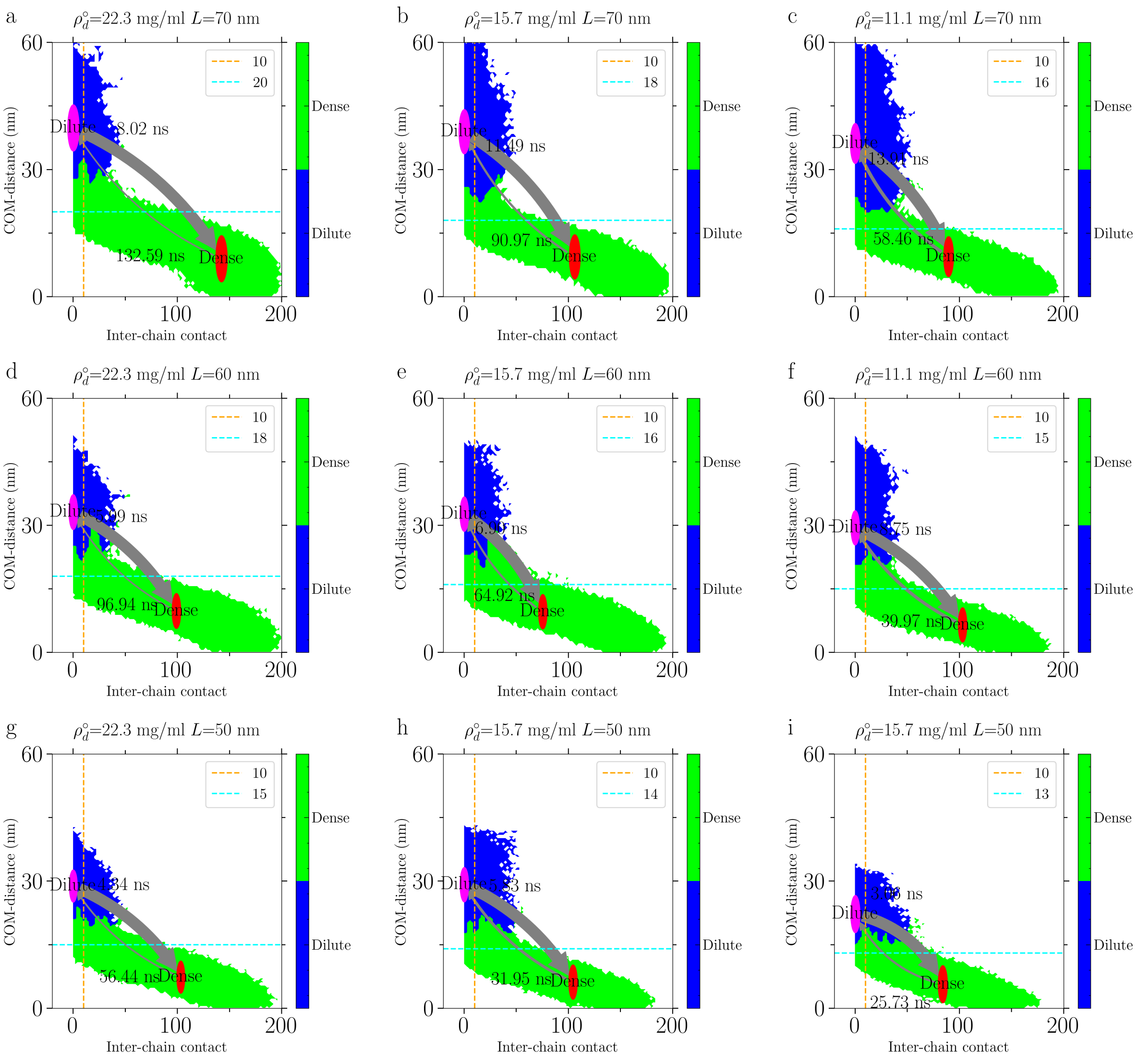


Figure S8: 2D MSM analysis for FUS-LC

#### Force-field parameters

The potential energy between two amino acids $i$ and $j$ at distance $r$ is given by:

$$V_{ij} = V_{b,ij} + V_{elec,ij} + V_{LJ,ij} \tag{3}$$

where $V_{b,ij}$ is a harmonic potential between bonded amino acids, with an equilibrium distance of 0.38 nm and a spring constant of 10 $kJ/mol/nm^2$.

$V_{elec,ij}$ is the Coulombic interaction with a Debye-Hückel screening:

$$V_{elec,ij} = \frac{q_i q_j}{4\pi D r} exp(-r/\kappa) \tag{4}$$

with $\kappa$ equal to 1 nm. $V_{LJ,ij}$ is the Lennard-Jones short-range potential:

$$V_{LJ,ij} = 4\lambda_{ij}\epsilon((\frac{\sigma_{ij}}{r})^{12} - (\frac{\sigma_{ij}}{r})^{6}) \tag{5}$$

where $\sigma_{i,j}$ and $\lambda_{i,j}$ are computed with the arithmetic combination from the parameters in Table S2, and $\epsilon$ is equal to 0.648 $kJ/mol$ (see Coexistence Simulations section in Supplementary Material). All non-bonded interactions were truncated with a cut-off of 4 nm.

Table S2 shows our CG parameters for running simulations.

Table S2: Parameters of the CG residues

| Residue | $m_i$[Da] | $q_i$[e] | $\sigma_i$[nm] | $\lambda_i$[-] |
|---|---|---|---|---|
| A | 71 | 0.00 | 0.504 | 1/3 |
| C | 103 | 0.00 | 0.548 | 1/3 |
| D | 115 | -1.00 | 0.558 | 1/3 |
| E | 129 | -1.00 | 0.592 | 1/3 |
| F | 147 | 0.00 | 0.636 | 1 |
| G | 57 | 0.00 | 0.45 | 1/3 |
| H | 137 | 0.50 | 0.608 | 1/3 |
| I | 113 | 0.00 | 0.618 | 1/3 |
| K | 128 | 1.00 | 0.636 | 1/3 |
| L | 113 | 0.00 | 0.618 | 1/3 |
| M | 131 | 0.00 | 0.618 | 1/3 |
| N | 114 | 0.00 | 0.568 | 1/3 |
| P | 97 | 0.00 | 0.556 | 1/3 |
| Q | 128 | 0.00 | 0.602 | 1 |
| R | 156 | 1.00 | 0.656 | 1 |
| S | 87 | 0.00 | 0.518 | 1/3 |
| T | 101 | 0.00 | 0.562 | 1/3 |
| V | 99 | 0.00 | 0.586 | 1/3 |
| W | 186 | 0.00 | 0.678 | 1 |
| Y | 163 | 0.00 | 0.646 | 1 |

### Effect of parameters for cluster analysis

We have also applied a different segment scheme and $R_0$ value for cluster analysis, to demonstrate that adjusting the algorithm parameter would not affect the results and hence the predicted thermodynamics and kinetics. The systems tested are NDDX4 at $\rho_d^{\circ}$=32.9 mg/ml and $L$=70 nm and FUS-LC at $\rho_d^{\circ}$=22.3 and $L$=70 nm. Figures S9 compares the clustersize distributions estimated using different $R_0$ and $n_{\text{beads}}$. The results conclude that the parameters used in cluster analysis do not affect the results.

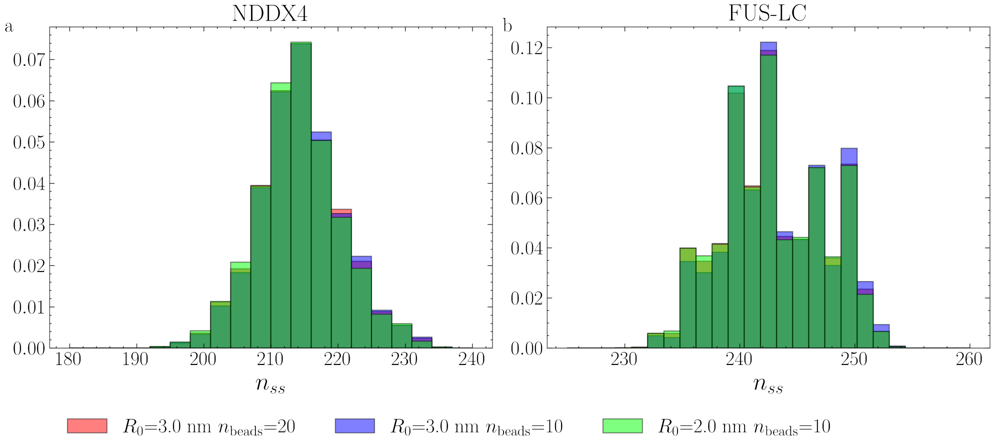


Figure S9: Clustersize distributions estimated using different $R_0$ and $n_{\text{beads}}$ for NDDX4 at $\rho_d^{\circ}$=32.9 mg/ml and $L$=70 nm and FUS-LC at $\rho_d^{\circ}$=22.3 and $L$=70 nm.

### Peptide inter-chain interaction

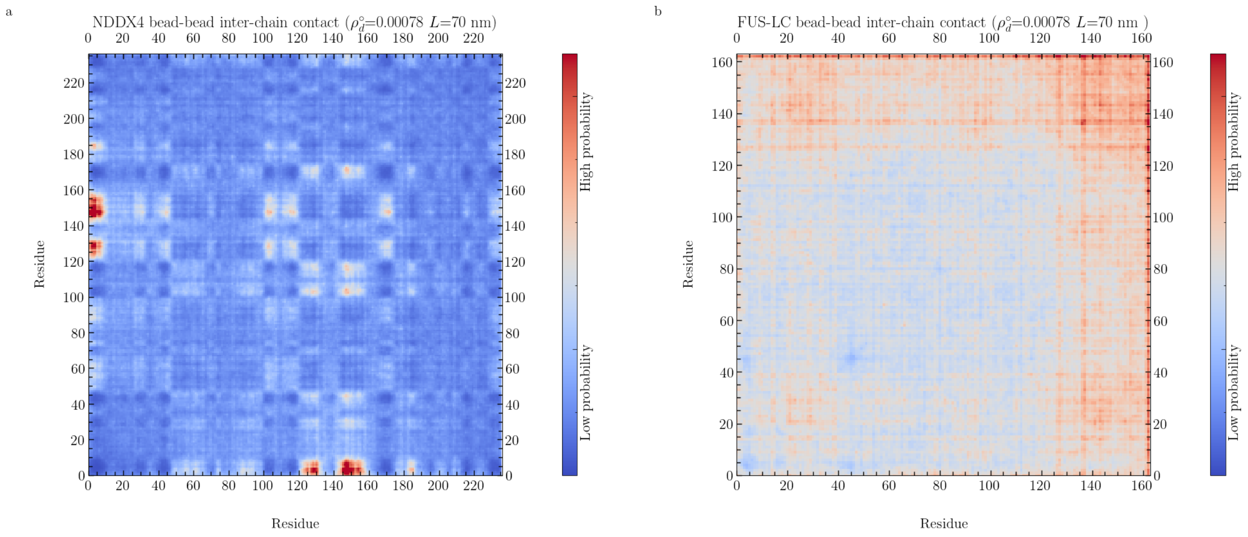


Figure S10: Bead-bead inter-chain contacts for NDDX4 and FUS-LC.

### Coexistence Simulations

We performed phase coexistence simulations [11,12] with the following protocol for both systems. 100 copies of the protein molecules were inserted in random positions and orientations in a $20 \times 20 \times 40$ nm$^3$ ($x \times y \times z$) box using the gmx insert-molecules tool, before extending the simulation box in the $z$ direction to a length of 200nm, keeping the protein center of mass in the center of the simulation box. Finally, we performed production simulations in the $NVT$ ensemble at a temperature of 300K using a leap-frog stochastic dynamics integrator with a timestep of 0.01 ps and a relaxation time of 25 ps. We evaluated the protein concentration along the $z$ axis of the box with the gmx density tool in Gromacs 2018.3, [13] centering the dense liquid protein phase (periodic in $x$ and $y$) at $z = 0$ at each frame. The dense phase concentration was determined from the values of density around $z = 0$, while dilute phase concentrations were evaluated by averaging concentration values at 60nm$\leq|z|\leq$100nm. We investigated three values of $\varepsilon$, namely 0.627, 0.648, and 0.670 kJ/mol, and chose a value of 0.648 kJ/mol as the one that best reproduced the equilibrium densities of both NDDX4 and FUS-LC (see Fig S11). The equilibrium densities of the condensed and dilute phases obtained from slab simulations are 335.93±10.60 and 6.97±0.30 mg/ml for NDDX4, 484.44±2.60 and 0.74±0.68 mg/ml for FUS-LC. The corresponding experimentally measured densities are: 380 mg/ml, and 7 mg/ml for NDDX4, [14] and 2 mg/ml [15] and 477 mg/ml [16] for FUS-LC.

The interfacial free energy from coexistence simulations was estimated following the expression reported in Tejedor, et al. [17] :

$$\gamma = \frac{L_N}{2}(p_N - p_T) \tag{6}$$

Where $L_N$ is the length of the simulation box in z direction, $p_N$ is the component of the pressure tensor normal to the slab surface ($p_{ZZ}$) and $p_T$ is the average of the components of the pressure tensor tangential to the slab surface ($p_{XX}$ and $p_{YY}$). The components of the pressure tensor were evaluated with the gmx energy tool available in Gromacs. [13] The surface tension estimates from the slab simulations are, for NDDX4 0.125±0.099 mN/m and for FUS-LC 0.291±0.026 mN/m.

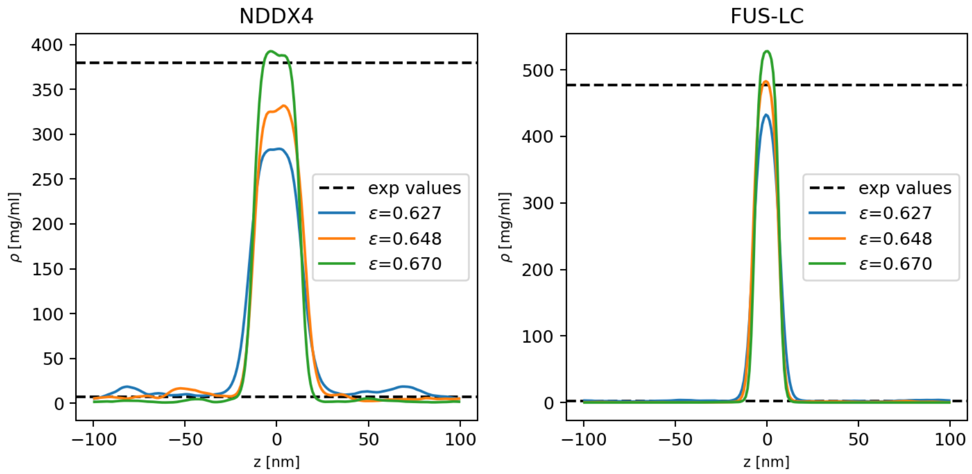


Figure S11: Density profiles from slab simulations of NDDX4 and FUS-LC for different values of the energy scale parameter $\varepsilon$.

(7) Vymětal, J.; Vondrášek, J. Gyration- and Inertia-Tensor-Based Collective Coordinates for Metadynam-

ics. Application on the Conformational Behavior of Polyalanine Peptides and Trp-Cage Folding. *The Journal of Physical Chemistry A* **2011**, *115*, 11455–11465.
